# Universal principles of lineage architecture and stem cell identity in renewing tissues

**DOI:** 10.1101/2020.03.10.984898

**Authors:** Philip Greulich, Ben D. MacArthur, Cristina Parigini, Rubén J. Sánchez-García

**Author notes:** P.G., B.D.M. and R.S.G conceived the project, P.G., B.D.M. and R.S.G wrote the paper, P.G. supervised the research, C.P. carried out the numerical analysis, and P.G. and R.S.G. carried out the mathematical analysis. Correspondence should be addressed to P.G.

## Abstract

Adult tissues in multicellular organisms typically contain a variety of stem, progenitor and differentiated cell types arranged in a lineage hierarchy that regulates healthy tissue turnover and repair. Lineage hierarchies in disparate tissues often exhibit common features, yet the general principles regulating their architecture are not known. Here, we provide a formal framework for understanding the relationship between cell molecular ‘states’ (patterns of gene, protein expression etc. in the cell) and cell ‘types’ that uses notions from network science to decompose the structure of cell state trajectories into functional units. Using this framework we show that many widely experimentally observed features of cell lineage architectures – including the fact that a single adult stem cell type always resides at the apex of a lineage hierarchy – arise as a natural consequence of homeostasis, and indeed are the only possible way that lineage architectures can be constructed to support homeostasis in renewing tissues. Furthermore, under suitable feedback regulation, for example from the stem cell niche, we show that the property of ‘stemness’ is entirely determined by the cell environment. Thus, we argue that stem cell identities are contextual and not determined by hard-wired, cell-intrinsic, characteristics.

## 1. Introduction

Adult stem cells were first identified in the 1960s context of hematopoesis (1, 2) and are now believed to reside in all renewing tissues. Because of their ubiquity and regenerative importance, the quest to find a complete set of defining characteristics of adult stem cells has been a longstanding aim of stem cell and developmental biology. However, despite numerous attempts, we still lack an unambiguous characterisation, and the stem cell concept is still widely disputed (3).

Historically, the stem cell identity has been viewed as a hard-wired phenotype, which is encoded in distinct patterns of gene and protein expression that confer to the cell the ability to maintain its own state while regenerating the cells of a whole tissue. Modern experimental methods have, however, cast doubt on this interpretation. For example, single-cell transcriptomics experiments have revealed a greater degree of molecular heterogeneity within apparently functionally ‘pure’ cell populations (including stem cell populations) than previously envisaged, and the search for unique stem cell markers has often, accordingly, proven challenging (4, 5). Perhaps more significantly, genetic cell lineage tracing studies have shown that many cells can exhibit a high degree of functional plasticity. For instance, cells which under homeostatic conditions are committed to differentiation are, in other contexts, such as regeneration or cancer, apparently able to acquire stem cell features, and can re-populate the stem cell pool via de-differentiation (6–11). In response to these studies it has been suggested that rather than being a hard-wired cell-intrinsic property, ‘stemness’ – i.e. the attribute of being a stem cell – is a functional, dynamical feature of cells that depends on the cellular environment (3, 12).

In practice, the most commonly accepted characterisation of stem cells is purely pragmatic: an adult stem cell is a cell which has the capability to (i) maintain its own population via self-renewing cell divisions and (ii) produce all cells of a specific tissue compartment via differentiation. According to this functional characterisation, ‘stemness’ is assessed by a cell’s lineage potential – i.e. what a stem cell and its progeny can become in the future, rather than by its molecular characteristics at present.

Here we will take this basic functional definition as a starting point and seek to examine the relationships between intracellular molecular ‘states’ and cell ‘types’ in order to clarify, precisely, what we mean when we refer to a stem cell. We start by acknowledging that a cell may be characterised by its internal molecular state (which we will define precisely shortly), and the possible changes in cell states which confer changes in cell function, for example when differentiating. It is reasonable, therefore, to define a cell both in terms of its current state, and the states that it may reach in the future under appropriate stimulus: that is, by the set of admissable cell state trajectories. Notably, such cell state trajectories may not just be simple paths: they can have a complex topology, including branching points at which cells make cell fate ‘decisions’ and closed loops or cycles (for example, the cell cycle).

Our essential insight is that we can represent all possible cell state trajectories as a directed network, in which nodes are cell states and links are admissible cell state transitions. Cell state trajectories are then directed paths on this network. Importantly, the structure of this network defines which trajectories are allowed and which are not and so is of central importance.

Here we analyse the general structural and dynamical properties of this cell state network, and use this analysis to precisely define stem cell identities and cell lineages in adult renewing tissues. Typically, we have very limited experimental information available about this potentially very complex network of cell states and transitions. However, we will show, using notions from dynamical systems and network theory, that under very mild and general assumptions concerning the underlying dynamics, the topology of an adult cell state network is necessarily very constrained. Specifically, the network can be naturally decomposed into smaller units, called strongly connected components (SCCs), which can be naturally associated with cell types. The network of possible transitions between SCCs then represents the lineage hierarchy of cell types in the tissue. Furthermore, if homeostasis is to be maintained, as is the case in most adult tissues, then the lineage hierarchy is additionally constrained.

The key finding of this article is that functional definitions of ‘stem’ and committed cells arise naturally within this framework. Particularly, there is an inherent relationship between the classification of stem versus committed cell types and their position in the cell lineage hierarchy: every cell lineage of a homeostatic renewing tissue must have a single stem cell type at its apex, and nowhere else. While this is commonly experimentally observed, there is currently no rationale for why this is the case. We demonstrate that this commonly observed structural hierarchy is not a mere coincidence of evolution, but instead, is the only possible lineage architecture that is able to maintain homeostasis in renewing tissues, and must therefore be present in any complex, multi-cellular, life form. Within this framework, it also becomes apparent that stemness is not necessarily a cell-intrinsic property, but rather is an emergent property of populations of cells interacting with their cellular environment, for example via crowding regulation or instruction from a niche.

## 2. Results

### A. Cell state networks

To start, we assume that a cell’s phenotype can be characterised by its internal molecular composition, including expression of RNAs, proteins, and metabolic components, etc. Here, we are interested not only in the current state of the cell, but also its potential, including its propensity to divide, change its phenotypic state, die or emigrate out of the tissue of interest. Rather than restrict to any specific technology (e.g. single cell RNA sequencing) we define a cell’s state in terms of the molecular features (measurable or not) which determine these propensities. While cell state is often described as a continuous quantity (13–16), here we adopt a discrete description with a finite number of states, which will allow us to use network theory for our analysis. Such a discretisation could be, for example, the commonly performed clustering of transcriptome data, or by simply acknowledging that molecule numbers are discrete. Our results, however, do not depend on the way the cell states are discretised, as we will discuss later.

To formalise our arguments, assume that there are ***m*** distinct cell states. Since we are interested in cell state trajectories broadly defined, we generally need to consider three cell processes: cell division, direct transitions between cell states (e.g. differentiation), and cell loss (by death or emigration). Consider a cell in state ***i*** = 1, 2, …, ***m***. We can represent these three cell processes schematically as

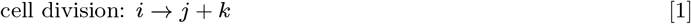

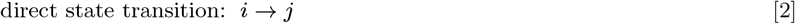

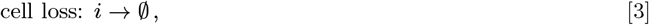

where the empty set symbol 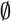 indicates ‘cell loss’ and ***j, k*** = 1, 2, …, *m* denote cell states that are possibly different from *i*. In particular, note that daughter cells from a cell division may or may not adopt states different to the mother cell’s (any two or all of the states ***i, j, k*** in Eq. (1) may be equal or not). The processes Eq. (1)–Eq. (3) determine the general dynamics of cells in a tissue. Equations describing the dynamics of the expected number of cells in each state can be easily derived from Eq. (1)–Eq. (3), and are provided in Box 1.

Notably, we can represent the cell dynamics Eq. (1)–Eq. (3) as a network of cell states, which we will refer to as the *cell state network*. The nodes of this network are the cell states ***i*** = 1, 2, …, ***m*** (according to our chosen representation) and directed links correspond to admissible cell state transitions: there is a directed link from node ***i*** to node ***j*** whenever a transition from cell state ***i*** to cell state ***j*** is possible via cell division or direct state transition. Directed paths (i.e. sequences of directed links) in this network represent all possible *cell state trajectories*.

#### Box 1: Dynamics of expected cell numbers

Let ***n_i_***(***t***) denote the expected number of cells in state ***i*** at time ***t***. Furthermore, let us write the rate of cell division in state ***i*** as ***λ_i_***, the rate of direct transition from state ***i*** to state ***j*** as ***ω_i→j_***, the rate of cell loss in state ***i*** as ***d_i_***, and the probability that a cell in state ***i*** will produce daughter cells in state ***j*** and ***k*** when it divides as 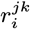. The time evolution of the expected number of cells of type ***i*** is given by

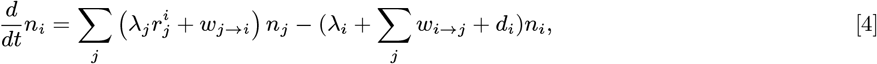

where 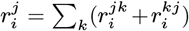 is the expected number of cells in state ***j*** produced upon division of a cell in state ***i***. We can write this system compactly in matrix form as follows. First, define the *total transition rate* from ***i*** to ***j*** as 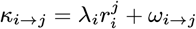, which combines all transitions from state ***i*** to state ***j***, via cell division Eq. (1) and direct state transitions Eq. (2). Note that ***κ_i→j_*** ≥ 0 for all ***i, j***. Also, define the *effective loss rate at **i*** as 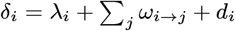, which combines transitions out of state *i* and cell loss. Using this notation, we can express the dynamical system, Eq. (4), in matrix form as

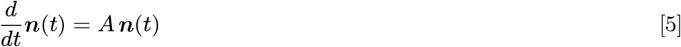

where ***A*** is the ***m*** × ***m*** matrix with (***i, j***)-entry ***κ_i→j_*** if ***i*** ≠ ***j*** and ***κ_i→i_*** − ***δ_i_*** if ***i*** = ***j***, and ***n***(***t***) = (***n*_1_**(***t***), ***n*_2_**(***t***), …, ***nm*** (***t***)).

The matrix ***A*** encodes the weighted cell state network described in the main text. This network contains all the information about the cell dynamics: the nodes represent the cell states ***i*** = 1, 2, …, ***m*** and any link from node (i.e. cell state) ***i*** to node ***j*** has weight ***κ_i→j_***, representing how likely this transition is. In particular, a link from cell state ***i*** to cell state ***j*** only exists if the transition is possible, that is, if ***κ_i→j_*** > 0. Note that self-loops (links from a node to itself) may occur, and that cell loss 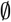 is not a node on the network, although we will add it to the graphical representation of our networks for clarity.

In general, the parameters ***λ_i_***, 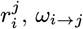, ***d_i_*** (***i, j*** = 1, …, ***m***) may depend on ***n*_1_, *n*_2_**, …, resulting in Eq. (4) becoming non-linear. This case is discussed further below, with details provided in Box 3. However, when the tissue is at homeostasis the expected cell numbers do not change under the dynamics (that is, mathematically, 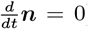). In this case, the dynamics in both the linear and nonlinear regimes are determined by the properties of the matrix ***A***, as we show in the main text.

### B. Cell types and strongly connected components

Cell types are typically defined by molecular characteristics, for example expression of surface markers or via clustering from single cell transcriptome data. Although pragmatic, such taxonomies do not provide a rationale for why certain molecular states are associated with specific cell functions. However, when viewed from the network perspective, we are able to classify cell states according to the role they play in tissue development and maintenance via their position in the cell state network.

To define a cell type in this context, we propose that cells of the same type should have the same lineage potential. Thus, any two states ***i*** and ***j*** belonging to the same cell type should share the same outgoing cell state trajectories. Mathematically, this is the case when states ***i*** and ***j*** are mutually reachable via cell state trajectories (directed paths in the cell state network). In this case, because cells can reversibly switch between states ***i*** and ***j***, any state reachable from ***i*** is also reachable from ***j***. In the terminology of network theory, two such states are said to be *strongly connected* (more precisely: nodes ***i*** and ***j*** of a directed network are strongly connected if there exist directed paths both from ***i*** to ***j***, and from ***j*** to ***i***, that is, ***i*** and ***j*** are connected by a cyclic trajectory). Extending this idea, it is natural to group states that are strongly connected with each other and we propose that each such grouping represents one cell type. Such a grouping of states (or, more generally, nodes of a directed network) is called a *strongly connected component* (SCC). Importantly, any directed network can be decomposed uniquely into SCCs (17), that is, the maximal groupings of mutually reachable nodes (see Fig. 1 for an example). From a biological viewpoint, it is reasonable to consider only *non-trivial* SCCs, i.e. those that contain at least one link, that is, either more than one node or one node with a self-link. These ideas motivate the following formal definition of a cell type in the context of development and tissue maintenance:

**Fig. 1.**
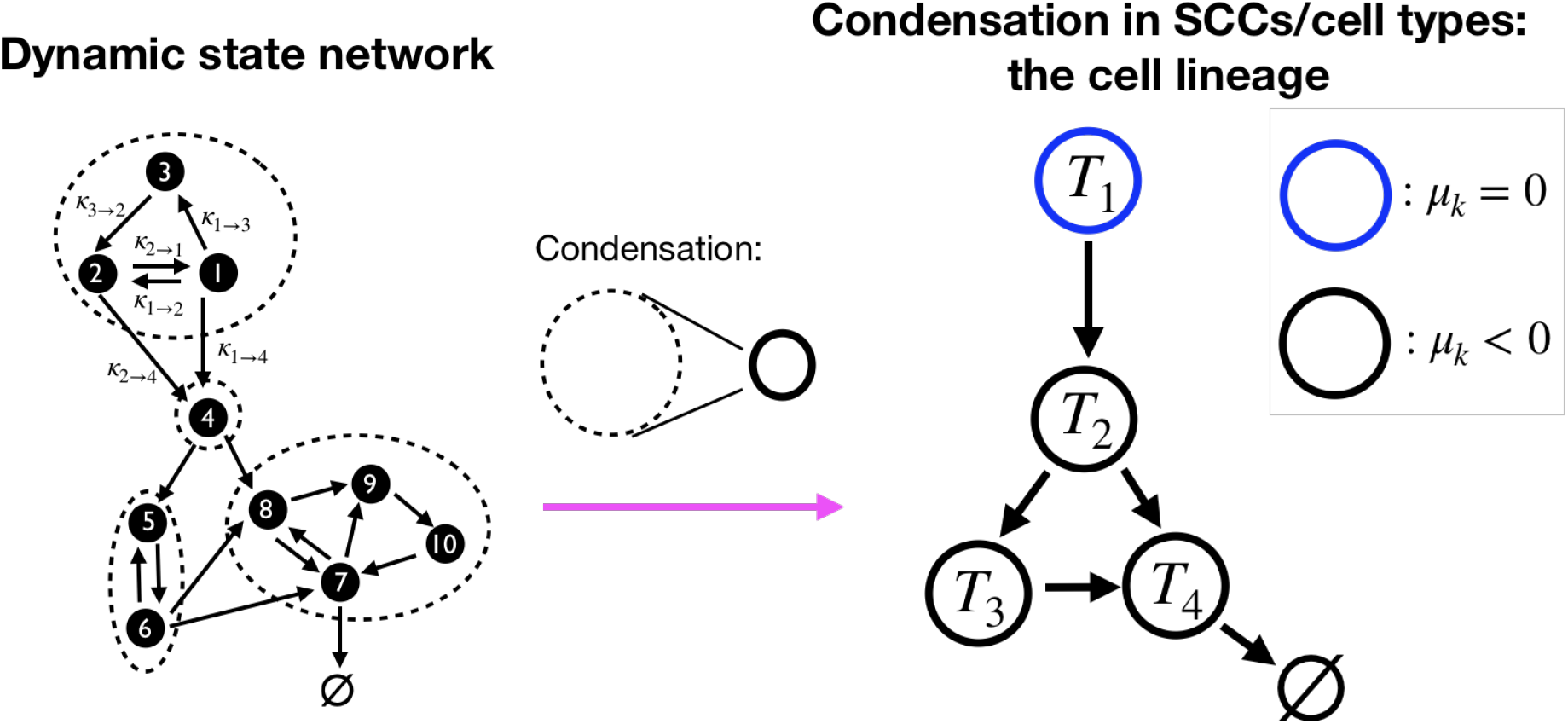
Grouping of cell states into cell types. (Left) A hypothetical cell state network. In this network intracellular molecular states are represented as nodes and possible transitions between states as links (to highlight the dynamics, some links are annotated with the total transition rates *κ_i→j_* as defined in Box 1). Cell loss (via death or emigration) is represented by the empty set symbol 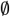. The dashed circles comprise states which are mutually reachable by directed paths in either direction, and thus are strongly connected components (SCCs) of the network. In our formulation, the SCCs represent cell types. (Right) When cell states are grouped into SCCs associated with cell types, the corresponding condensation forms a directed network without cycles, that encodes possible transitions between cell types. This network has a natural hierarchical structure: it admits an ordering of the nodes (cell types) *T*_1_, *T*_2_, … such that all transitions respect the ordering, that is, if there is a link, or a trajectory, from *T_k_* to *T_l_* then *k* ≤ *l*. Here, we show one such hierarchical ordering (*T*_1_ to *T*_4_). Additionally, we show a possible distribution of transient (black circles) and self-renewing (blue circles) cell types *T_k_*, *k* = 1, …, 4, depending on their growth parameter *μ_k_* (see Box 2). Finally, note that different choices of cell states will result in a different cell state network (left) but the same condensation network of cell types (right) —see main text.

#### Definition 1.

***A cell type** is a non-trivial strongly connected component of a cell state network*.

Notably, this definition is independent of discretisation of cell state trajectories: non-trivial SCCs are composed of nodes connected by cyclic trajectories, and the latter does not change upon a finer discretisation of cell state trajectories.

In the following sections, we will show that this simple definition of cell type naturally implies a number of well-known concepts and principles from stem cell and developmental biology, and, in particular, leads to a simple and unique definition of an adult stem cell.

### C. Condensation of the cell state network

To better understand the structure of the cell state network it is convenient to consider relationships between cell types (SCCs). To do so we construct a network within which cell types are nodes, and there is a directed link from cell type ***A*** to cell type ***B*** (not equal to ***A***) whenever there is at least one link from a state of type ***A*** to a cell state of type ***B*** (see Fig. 1 for an example). An elementary result of graph theory states that the resulting network, which is known as the *condensation*, does not contain cycles (directed paths from a node to itself) and is, therefore, hierarchical (18). More precisely, we can order the cell types ***T*_1_**, ***T*_2_**, …, such that if there is a trajectory (i.e. a directed path) from ***T_k_*** to ***T_l_*** then ***k*** ≤ ***l*** (Fig. 1). In this case, we will say that ***T_k_*** is *upstream* of ***T_l_*** and ***T_l_*** is *downstream* of ***T_k_***. Biologically, this means that cell types are necessarily ordered in a hierarchy, as is commonly observed.

This hierarchy is useful because it encodes the essential lineage structure of the tissue under consideration (we will consider some general structural features of this hierarchy shortly). However, the structure of this hierarchy does not, alone, determine the proliferative dynamics of the tissue. For that purpose, we also need to assess the intrinsic proliferative properties of each of the cell types present (i.e. its ability to perpetuate its own numbers). In Box 2, we show that the intrinsic dynamics of the ***k***th cell type, ***T_k_***, can be uniquely characterised by a single number, its growth parameter ***μ_k_***, which determines the long-term dynamics of cells of that type, when transitions from/to other cell types are neglected. In particular, if ***μ_k_*** > 0 then the expected number of cells of type ***T_k_*** increases without bound; if ***μ_k_*** < 0 then the expected number of cells of type ***T_k_*** decreases until the pool vanishes; and if ***μ_k_*** = 0 then the expected number of cells of type ***T_k_*** stays constant (i.e. cells of that type maintain themselves at homeostasis) (19). This classification of cell types according to their intrinsic dynamics motivates the following definition:

#### Definition 2.

*We call a cell type **T** with growth parameter **μ**:*

1. transient *if **μ*** < 0,
2. hyper-proliferating *if **μ*** > 0,
3. self-renewing *if **μ** = 0*.

These three classes represent three distinctly different kinds of dynamics. Firstly, transient cell types will eventually vanish, unless these cells are replenished through incoming trajectories from other upstream cell types. This means that all transient cells and their progeny will eventually differentiate into another type or will be lost from the tissue (e.g. via cell death or emigration). Secondly, hyper-proliferating cell types will continually increase in number in the tissue. Although possible, this situation is clearly not physiological, yet may be encountered in certain pathologies, such as cancer. Thirdly, self-renewing cell types will maintain their numbers, on average, over time. In principle this definition allows for inert cells that perform no function in the tissue (i.e. do not divide, die, emigrate or exhibit any other behaviour that would affect their numbers). However, that situation only occurs in cell lineages that are not renewing, which we do not consider here. In renewing cell lineages, such cells must balance proliferation and loss and so exhibit the biological property of self-renewal that is characteristic of stem cells.

Collectively, these results indicate that important features of cell lineage architectures, such as the presence of self-renewing populations, and hierarchical ordering of cell types in a lineage tree, emerge naturally via consideration of general structural properties of the cell state network.

#### Box 2: Cell type dynamics

Consider a cell type ***T_k_*** consisting of states ***i*_1_, *i*_2_**, …, ***i_m_k__*** in the cell state network. The intrinsic dynamics of the cell type ***T_k_***, i.e. when neglecting transitions from/to other cell types, are determined by the equation

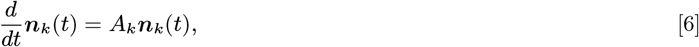

where ***n_k_***(***t***) = (***n*_*i*_1__**(***t***), ***n*_*i*_2__**(***t***), …, ***n_i_m_k___***(***t***)) is the vector of expected cell numbers at time ***t*** in the cell states ***i*_1_, *i*_2_**, …, ***i_m_k__*** comprising the cell type ***T_k_***. The matrix ***A_k_*** is the ***m_k_*** × ***m_k_*** submatrix of ***A*** (see Box 1) corresponding to the rows and columns ***i*_1_, *i*_2_**, …, ***i_m_k__***. The matrix ***A_k_*** is irreducible, that is, is the adjacency matrix of a strongly connected network (the SCC corresponding to ***T_k_***). Therefore, the Perron-Frobenius theorem guarantees that ***A_k_*** has a unique real maximal eigenvalue ***μ_k_***(20, 21). It is a basic result of dynamical systems theory that ***μ_k_*** determines the long-term dynamics of the system, and in particular the sign of ***μ_k_*** determines if the population will grow or decline with time (19) (see Section 1 of the Supplementary Material for further details). We call ***μ_k_*** the *growth parameter* of cell type ***T_k_***.

### D. Homeostasis imposes fundamental constraints on cell lineage architectures

So far we have considered the structure of the cell state and cell type networks without imposing any restriction on tissue proliferative dynamics. However, under normal circumstances, most adult tissues are maintained in a state of proliferative homeostasis. From a mathematical perspective, homeostasis represents a stable state of the dynamics in which the number of cells of each type in the tissue stays constant on average.

There are a number of ways that homeostasis can be maintained. In principle, a homogeneous cell population that consists of a self-renewing population of a single type, as defined above, would be by definition homeostatic. However, this is a particular situation that is not reflective of the complexity of most tissues. Typically, a tissue will consist of a range of different cells, each with different proliferative capacities, related in a complex hierarchy. Under these circumstances, for the cell population as a whole to be homeostatic, further conditions on the architecture of the lineage hierarchy need to be satisfied, which are discussed next.

The cell state dynamics described in Box 1 are an instance of a particular type of dynamical system called a *linear cooperative system* (a linear system where all couplings between distinct states are non-negative; in our case, these couplings are *κ_i→j_* ≥ 0, see Box 1). Because we wish to examine dynamics close to homeostatic equilibrium, in this section we will assume that the model parameters (*κ_i→j_* and *δ_i_*, see Box 1) are all constant. By so doing we are assuming that, close to equilibrium, the non-linear dynamics may be approximated by the linear system described in Box 1. Generalisations to include non-linear environmental effects, specifically crowding feedback, will be discussed in the next section.

We have recently derived general mathematical conditions that ensure stability of homeostasis in linear cooperative systems (22). These purely mathematical conditions have direct biological implications, which are outlined in Section 1 of the Supplemental Material. In particular, we show that homeostasis can be maintained in a renewing tissue if and only if:

1. No hyper-proliferating cell type is present.
2. At least one self-renewing cell type is present.
3. There are no cell state trajectories from one self-renewing cell type to another.

The third of these conditions has an important implication: since trajectories between self-renewing cell types cannot occur, one cannot be upstream of another. In combination with condition 2, this implies that each adult homeostatic lineage hierarchy must have a self-renewing cell type at its apex and nowhere else. Nonetheless, cell types upstream of a self-renewing type may exist transiently, in non-homeostatic situations as during development.

With these insights, we can classify each cell type according to its growth parameter and relative position in the hierarchy into one of the following three classes.

*Developing cells (D)*: These are transient cells (growth parameter ***μ*** < 0) upstream of a self-renewing cell type in the lineage hierarchy. Developing cells may transiently divide and differentiate but, since they are not replenished by a self-renewing cell type, they disappear as homeostasis is approached and are thus not present in adult, homeostatic tissues. We therefore associate these cells with developmental populations that do not survive into adulthood (e.g. embryonic stem cells).
*Committed cells (C)*: These are transient cells (growth parameter ***μ*** < 0) downstream of a self-renewing cell type. They may divide, and all their progeny are eventually lost (e.g. to death or emigration). However, this population does not disappear since it is maintained at homeostasis by replenishment from an upstream self-renewing population. We associate committed cells with adult somatic cells that are not stem cells.
*Adult stem cells (S)*: These are self-renewing cells (growth parameter ***μ*** = 0, maintaining their population) which can differentiate into at least one committed cell type downstream.

From the structural conditions above and these definitions we arrive at the following fundamental principle for cellular hierarchies:

#### Principle 1.

*Adult cell lineage hierarchies of renewing tissues must have a stem cell type at their apex and, conversely, any stem cell type must be at the apex of a cell lineage hierarchy*.

Note that multiple apexes of a lineage can in principle exist (see Fig. 2), as is for instance conjectured for mouse mammary epithelium lineage (23).

**Fig. 2.**
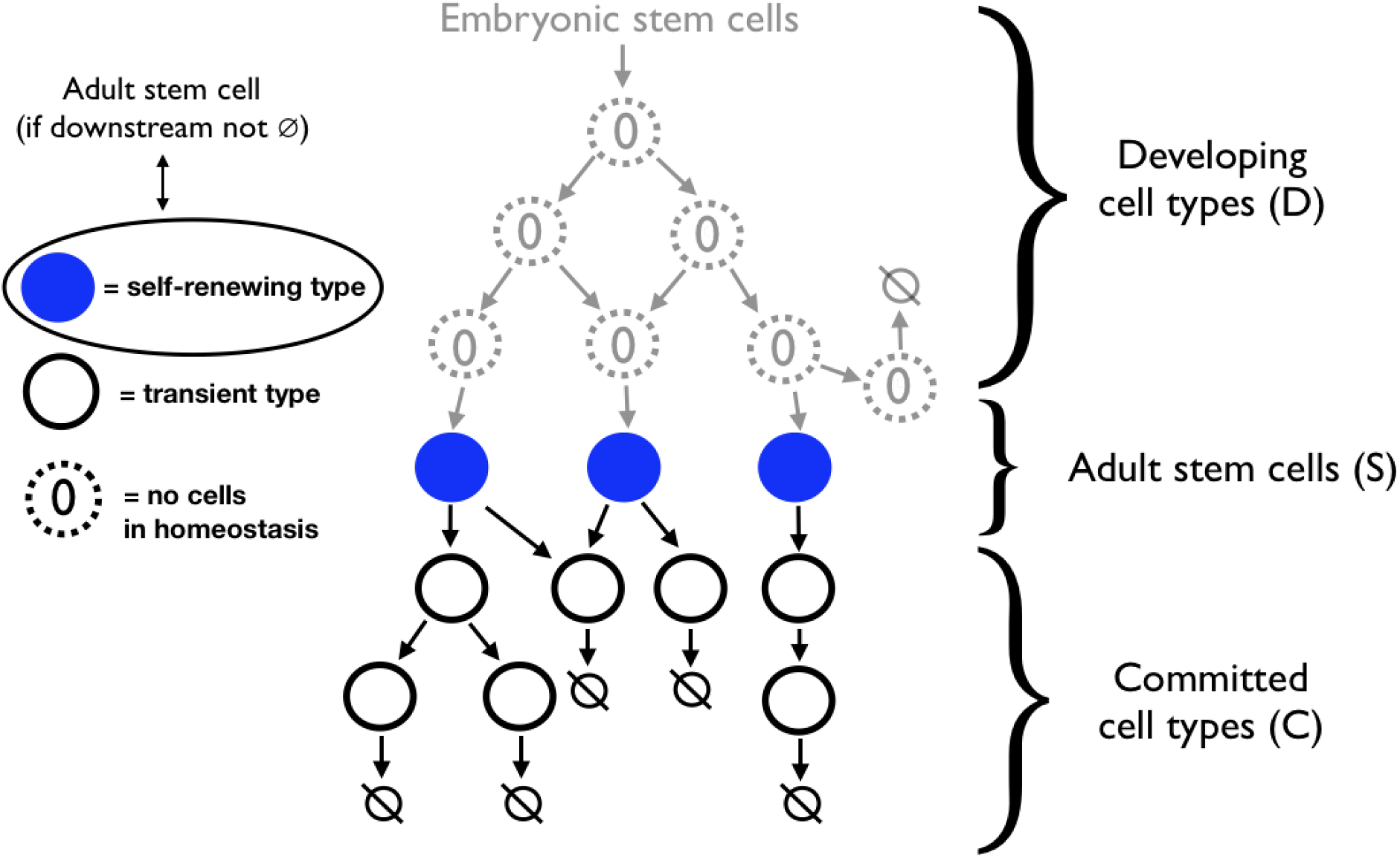
Illustration of a typical homeostatic cell lineage. Each circle represents a cell type (see Fig. 1), which comprises a maximal set of mutually reachable cell states. Faint grey dashed circles are developing cell types (D) which lead into a self-renewing state, but which disappear in the adult tissue. Filled-in (blue) circles represent self-renewing cell types. Black bold circles are committed cell types (C) which can differentiate and whose progeny is eventually lost. Crucially, along each lineage trajectory (a path on the network) only a single self-renewing cell type can exist, which we identify as adult stem cells (S). Thus, in the long term, a single stem cell type must be at each apex of the homeostatic lineage.

This general principle and the conditions above have a number of implications. For example, it immediately follows that only three patterns of stem cell division are allowed:

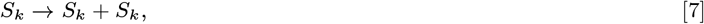

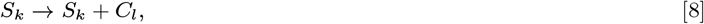

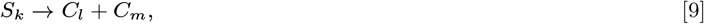

where ***S_k_*** is an adult stem cell type of type ***k***, and ***C_l_***, ***C_m_*** are (possibly different) committed cell types. These three division types correspond precisely to what is known experimentally about stem cell divisions. In particular, stem cells may divide in the following three ways: (1) symmetric self-renewal divisions (***S* → *S* + *S***), (2) asymmetric self-renewal divisions (***S* → *S* + *C***) and (3) differentiation division (***S* → *C* + *C***). However, following the conditions for homeostasis above, we can say more. For instance, a stem cell division producing one or more stem cells of different types, i.e. a division of the form ***S_k_* → *S_l_* + *S_m_*** or ***S_k_* → *S_l_* + *C*** in which ***l* = *k*** or ***m* = *k*** is not possible, as it would contradict condition 3 above. Furthermore, stem cell divisions that produce developing cells, and divisions of a committed cell to produce a stem cell, are also not possible, according to Principle 1. To make these ideas more concrete, examples of lineage hierarchies that do/do not fulfill these principles are shown in Figures 2 and 3.

Collectively, these results show that characteristic properties of lineage architectures may be derived from first principles via consideration of the relationship between cell states and cell types, and from dynamical constraints under homeostasis.

**Fig. 3.**
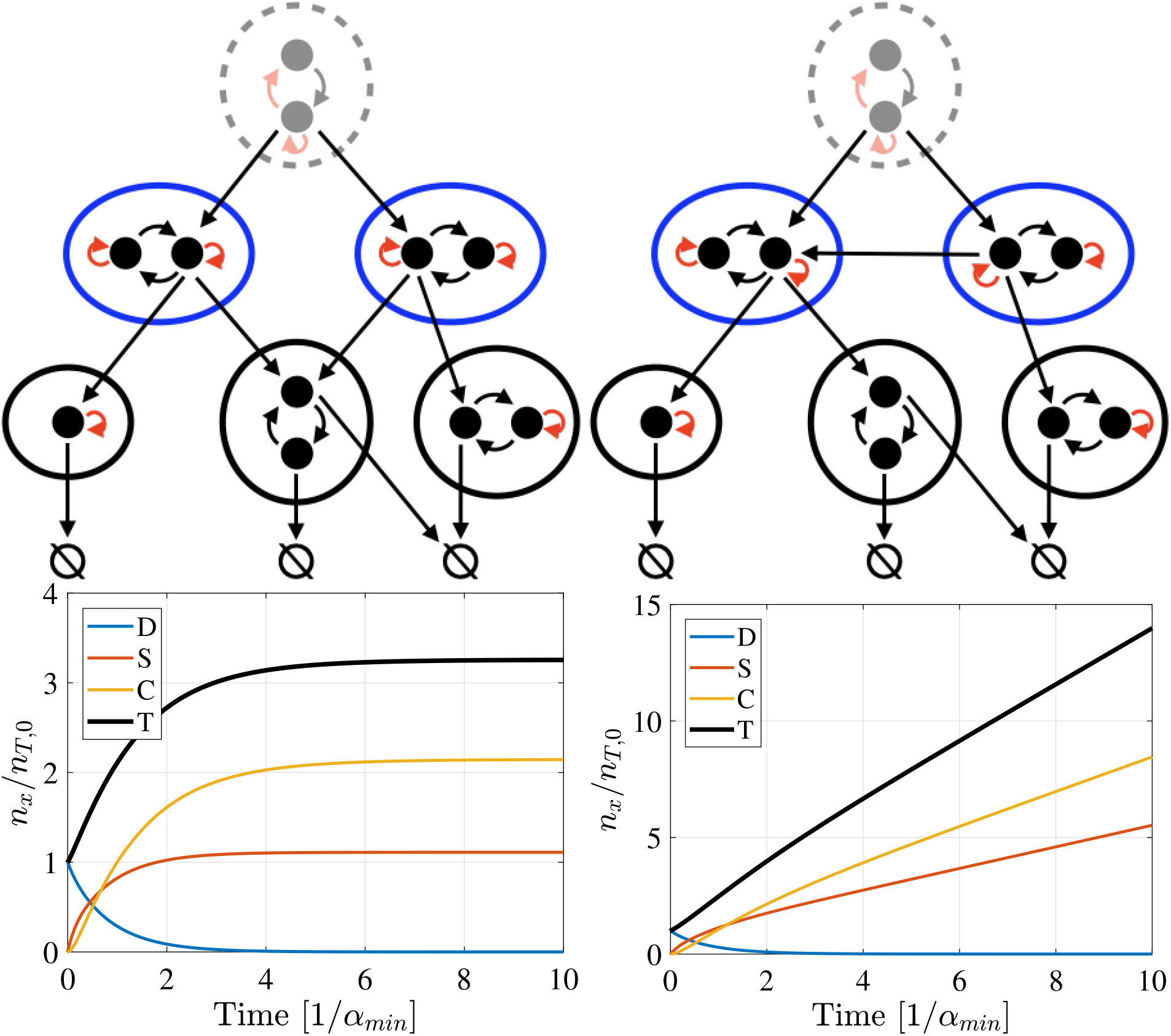
Lineage hierarchy and cell dynamics. (Top) Two cell state networks: black dots represent cell states, black arrows represent direct cell state transitions (according to Eq. (2)), red arrows represent transitions through cell divisions (according to Eq. (1)), and 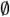 denotes cell loss (via cell death or emigration, for example). We draw circles around cell states of the same type (the SCCs of the underlying cell state network): black circles represent transient cell types, and blue circles self-renewing cell types. Grey faint states and circles denote developing cell states and types. (Bottom) Simulations of expected cell numbers obtained by solving Eq. (4)numerically for the networks immediately above. (Left panels) This network (top left) fulfills the criteria for homeostasis, and thus (bottom left) the expected numbers of stem cells (*n_S_*), and committed cells (*n_C_*) reach a steady state, while the expected number of developing cells (*n_D_*) tends to zero. The total expected number of cells, *n_T_*, also reaches a steady state. (Right panels) This network (top right) does not fulfill the criteria for homeostasis since two self-renewing cell types are connected by a direct link and thus (bottom right) the cell numbers *n_S_* and *n_C_* diverge. Simulations are numerical solutions of the dynamical equation Eq. (4), which are shown as function of time (scaled by the inverse of smallest rate *α_min_* = min_*i,j*_(*λ_i_*, *ω_i→j_*, *d_i_*)).

### E. Environmental regulation of the stem cell identity

In order to be self-renewing, the parameters of a stem cell type (rates of division, cell state transitions, and cell loss) need to be finely tuned to achieve a growth parameter of exactly zero, (***μ*** = 0, see Box 2). Any slight deviation from this value will lead to a loss of the self-renewing capacity and thus of the stem cell phenotype. In the absence of a homeostatic control mechanism such fine tuning is biologically implausible. In reality, cells are embedded in a tissue environment and may respond to signalling cues from other cells. An simple example of such regulation is crowding feedback, in which cells sense the density of cells in their local micro-environment, for instance via mechanosensing (24–26) or through the limited availability of growth signalling factors (27), and respond by adjusting their propensity to divide.

We can incorporate such regulation in our framework by assuming that the kinetic parameters associated with the dynamics in Eq. (1)-Eq. (3), as defined in Box 1, are not constant, but depend on the cell number density. A kinetic parameter may increase with the cell density (i.e. exhibit *positive crowding dependence*) or decrease with it (exhibit *negative crowding dependence*), or neither. In Box 3 and Section 2 of the Supplemental Material, we show that a simple sign criterion for the crowding dependence of the kinetic parameters guarantees that the cell type at the apex of the cell lineage will acquire and maintain self-renewal. It follows from this condition that if all other cell types are transient, then the tissue as a whole will become homeostatic when this sign condition is fulfilled.

This feedback-based mechanism for regulating stemness is simple and robust against noise, as it does not depend on the exact values of the dynamic parameters, but only on the sign of their dependencies. Crucially, this means that self-renewal need not be a property intrinsically determined by a cell, but it may be acquired by any cell type that is at the apex of a lineage hierarchy, through interaction with the cellular environment.

#### Box 3: Regulation of self-renewal by crowding feedback

Consider a cell type ***T_a_*** at the apex of a cell lineage, which is not a priori a self-renewing type. Assume that the kinetic parameters of the cell dynamics, ***λ_i_***, ***ω_i→j_***, and ***d_i_*** (see Box 1), may depend on the density of cells in the ‘niche’, which we assume to be constituted by the cells of type ***T_a_***. For constant volume, the density is proportional to the number of cells of type ***T_a_***, 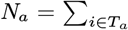, and thus these parameters are functions of ***N_a_***: ***λ_i_***(***N_a_***), ***ω_i→j_***(***N_a_***), and ***d_i_***(***N_a_***). The time evolution for cell numbers per state in ***T_a_*** is, according to Eq. (6), 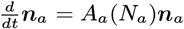. We note that, since the kinetic parameters are functions of ***N_a_***, so are the entries of the matrix ***A_a_*** and thus the growth parameter ***μ_a_*** = ***μ_a_***(***N_a_***) as well.

Now, for notational convenience, let us rename all kinetic parameters by ***α_j_***, numbered by 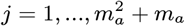, in the order ***λ*_1_, *λ*_2_**, …, ***ω*_1→2_, *ω*_1→3_, …, *d*_1_, *d*_2_**, In Section 2 of the Supplemental Material, we show that a sufficient condition for ***T_a_*** to attain self-renewing is

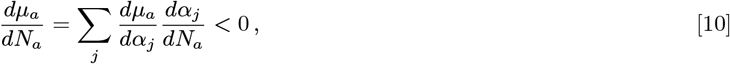

for all ***N_a_***. Recall that the crowding dependence of a parameter ***α*** is positive if ***dα/dN_a_*** > 0 and negative if ***dα/dN_a_*** < 0. For a cell type at the apex of a non-trivial lineage hierarchy the following criterion holds:

**Sign criterion for regulation of self-renewal:** If for all dynamic parameters ***α_j_***, the crowding dependence is positive (respectively negative), while the growth parameter dependence on ***α_j_*** is negative (respectively positive), then a cell type at the apex of a lineage hierarchy acquires and maintains self-renewal.

Because ***T_a_*** sits at the apex of a non-trivial lineage hierarchy (i.e. cells in ***T_a_*** have the capacity to differentiate into at least one other cell type), this means that the cells in ***T_a_*** gain stem cell status. Note that it is not required that all parameters ***α_j_*** have a suitable crowding dependence; in particular, for any parameter with ***dμ_a_/dα_j_*** = 0, the crowding dependence of ***α_j_*** is not relevant. An illustrative example of this result is shown in the Supplementary Material, Figure 1.

## 3. Discussion

Stem cells are fundamental to healthy tissue turnover and repair and have accordingly been the subject of concerted research efforts. However, despite more than 60 years of research, our understanding of the general principles that underpin stem cell dynamics remains incomplete. Here, we have proposed a new way to approach understanding stem cell identities from first principles based upon consideration of the relationships between cell states and cell types and general properties of proliferation in hierarchically structured populations. The resulting organisational principles are consistent with common conceptualisations of adult stem cells and lineage architectures, even though they emerge from consideration of the dynamical properties of cell lineage architectures at homeostasis rather than from explicitly biological considerations. Furthermore, our analysis indicates that these common properties are not coincidental but rather reflect fundamental and universal constraints on tissue dynamics at homeostasis. The common observation that stem cells sit at the apex of lineage hierarchies, for example, is not simply due to biological contingency: it is the only possible way that lineage architectures can be constructed to support homeostasis in renewing tissues.

It is important to note that our results only hold at homeostasis. In challenged conditions, such as wounding and cancer, lineage architectures may temporarily change to allow de-differentiation, for example. In such situations properties of the lineage hierarchy not accounted for in our formulation may also be important. For example, our Principle 1 does not exclude an additional pool of dormant stem cells residing in the lineage hierarchy above an adult stem cell population. Such dormant stem cells can not participate in homeostatic tissue renewal, and so play no role in healthy tissue turnover, but may nevertheless be present and become activated under conditions of stress.

Our results accord well with current experimental evidence in numerous tissues. For example, in most mammalian renewing epithelia, such as epidermis (28, 29), oesophagus (30, 31), and intestine (32), a single stem cell type, at the apex of the lineage hierarchy, can be identified. Other tissues, such as mammary gland, may exhibit multiple stem cell types in the same tissue. However, each of them resides on a separate apex of the cell lineage (23) and thus are consistent with the possible lineage architectures predicted here.

While most systems appear to have a single stem cell at the apex of each lineage hierarchy, the hematopoietic system is an interesting potential counter-example. Hematopoietic stem cells (HSCs) are perhaps the most well studied adult stem cells. In this system, two distinct types of stem cell are known to exist (along with numerous different kinds of progenitor cells) – long-term (LT) and short-term (ST) repopulating HSCs – and it has been suggested that they stand in a hierarchical relationship to one another (33). If so, this would violate Principle 1. While the functional distinction between LT-HSCs and ST-HSCs has not yet been fully characterised, there are several possibilities that would resolve this apparent contradiction. Firstly, it is notable that much of our understanding of HSC function comes from bone marrow transplantation experiments, which are fundamentally out of equilibrium and so outside the scope of our theory. Indeed, there is increasing evidence that the mechanisms of HSC proliferation in times of stress is fundamentally different to that under steady state. With this in mind it is possible that: (1) LT-HSCs constitute a dormant population that does not participate in normal homeostasis, but are activated when homeostasis is disturbed, e.g. after blood loss; (2) the ST-HSC population is not genuinely self-renewing, but gradually loses its proliferative potential on a long time scale (for instance, during aging). In our formulation this would mean that the ST-HSC growth parameter *μ* = −*ϵ* where 0 < *ϵ* ≪ 1; (3) LT-HSCs and ST-HSCs can inter-convert (even if only rarely). If so they would, according to our definition, effectively be a single stem cell type. These issues are hard to resolve experimentally, yet theoretical considerations can be a guide. It is likely that similar issues will arise as we gain deeper understanding of other systems. If so, then theoretical notions such as the ones we propose could be invaluable. We anticipate that future developments in stem cell biology will benefit from closer integration of experiment and theory which will yield a deeper understanding of the organisational principles that regulate stem cell dynamics.

## Supporting information

Supplemental Materials: Mathematical Derivations

## References

1. Siminovitch L, McCulloch EA, Till JE (1963) The Distribution of Colony-Forming Cells Among Spleen Colonies. Journal of cellular physiology 62:327.

2. Becker AJ, McCulloch EA, Till JE (1963) Cytological demonstration of the clonal nature of spleen colonies derived from transplanted mouse marrow cells. Nature 197:452.

3. Clevers H (2015) What is an adult stem cell? Science 350:4.

4. Ghadially R (2012) 25 years of epidermal stem cell research. Journal of Investigative Dermatology 132:797.

5. Alcolea MP (2017) Oesophageal Stem Cells and Cancer, in Stem Cell Microenvironments and Beyond, ed. Birbrair A. (Springer International Publishing, Cham), pp. 187.

6. Donati G, Watt FM (2015) Stem Cell Heterogeneity and Plasticity in Epithelia. Cell Stem Cell 16:465.

7. Donati G, et al. (2017) Wounding induces dedifferentiation of epidermal Gata6+ cells and acquisition of stem cell properties. Nature Cell Biology 19:603.

8. Tetteh PW, Farin HF, Clevers H (2015) Plasticity within stem cell hierarchies in mammalian epithelia. Trends in Cell Biology 25:100.

9. Tetteh PW, et al. (2016) Replacement of Lost Lgr5-Positive Stem Cells through Plasticity of Their Enterocyte-Lineage Daughters. Cell Stem Cell 18:203.

10. Koren S, Bentires-Alj M (2015) Review Breast Tumor Heterogeneity: Source of Fitness, Hurdle for Therapy. Molecular Cell 60:537.

11. Van Keymeulen A, et al. (2015) Reactivation of multipotency by oncogenic PIK3CA induces breast tumour heterogeneity. Nature 525:119.

12. Clevers H, Watt FM (2018) Defining adult stem cells by function, not by phenotype. Annual review of biochemistry 87:1015.

13. Weinreb C, Wolock S, Tusi BK, Socolovsky M, Klein AM (2018) Fundamental limits on dynamic inference from single-cell snapshots. Proc. Natl. Acad. Sci. 115:E2467.

14. Huang S (2012) The molecular and mathematical basis of Waddington’s epigenetic landscape: A framework for post-Darwinian biology? BioEssays 34:149.

15. Huang S (2010) Cell lineage determination in state space: A systems view brings flexibility to dogmatic canonical rules. PLoS Biology 8:8.

16. Trapnell C (2015) Defining cell types and states with single-cell genomics. Genome Research 25:1491.w

17. Bollobás B (2013) Modern graph theory. (Springer Science & Business Media).

18. Cormen TH (2009) Introduction to Algorithms. (MIT Press).

19. Strogatz S (1994) Nonlinear dynamics and chaos: with applications to physics, biology, chemistry, engineering. (CRC Press)

20. MacCluer CR (2000) The Many Proofs and Applications of Perron’s Theorem. SIAM Review 42:487.

21. Arrow KJ (1989) A “dynamic” proof of the Frobenuis-Perron theorem for Metzler matrices, in Probability, Statistics, and Mathematics, ed. Anderson TW, Athreya KB, Iglehart DL. (Academic Press) pp. 17.

22. Greulich P, MacArthur BD, Parigini C, Sánchez García RJ (2019) Stability and steady state of complex cooperative systems: a diakoptic approach. Royal Society Open Science 6:191090.

23. Pal B, et al. (2017) Construction of developmental lineage relationships in the mouse mammary gland by single-cell RNA profiling. Nature Communications 8:1627.

24. Puliafito A, et al. (2012) Collective and single cell behavior in epithelial contact inhibition. Proc. Natl. Acad. Sci. 109:739.

25. Eisenhoffer GT, Rosenblatt J (2013) Bringing balance by force: live cell extrusion controls epithelial cell numbers. Trends in cell biology 23:185.

26. Shraiman BI (2005) Mechanical feedback as a possible regulator of tissue growth. Proc. Natl. Acad. Sci. 102:3318.

27. Kitadate Y, et al. (2019) Competition for Mitogens Regulates Spermatogenic Stem Cell Homeostasis in an Open Niche. Cell Stem Cell 24:79.

28. Alonso L, Fuchs E (2003) Stem cells of the skin epithelium. Proc. Natl. Acad. Sci. 100:11830.

29. Clayton E, et al. (2007) A single type of progenitor cell maintains normal epidermis. Nature 446:185.

30. Seery JP, Watt FM (2000) Asymmetric stem-cell divisions define the architecture of human oesophageal epithelium. Current Biology 10:1447.

31. Seery JP (2002) Stem cells of the oesophageal epithelium. Journal of Cell Science 115:1783.

32. Barker N, et al. (2007) Identification of stem cells in small intestine and colon by marker gene Lgr5. Nature 449:1003.

33. Cheng H, Zheng Z, Cheng T (2019) New paradigms on hematopoietic stem cell differentiation. Protein and Cell 11:34.

