## Supplemental Materials: Mathematical Derivations for "Universal principles of lineage architecture and stem cell identity in renewing tissues"

#### 1 Conditions for homeostasis on lineage hierarchies

Here we adapt the generic conditions for the existence of a stable steady state in linear cooperative systems in Ref. [1] to the biological context in order to derive the conditions for homeostatic tissue cell dynamics given in the Main Text.

First, we note that the the generic cell dynamics studied in the Main Text (Eq. **[5]** in Box 1) represent a *linear cooperative system* (LCS). An LCS is a system of linear ordinary differential equations for functions of time $t$, $x_1(t), x_2(t), ..., x_m(t)$ which can be written in the form $\frac{d}{dt}\boldsymbol{x} = A\boldsymbol{x}$, where $\boldsymbol{x} = (x_1(t), x_2(t), ..., x_m(t))$ and $A$ is a constant matrix with non-negative off-diagonal elements, $a_{ij} \geqslant 0$ for $i \neq j$ [2]. These properties are fulfilled for Eq. **[5]**, Main Text, since the off-diagonal elements of $A$, the total transitions rates $\kappa_{i \to j} \geqslant 0$, are non-negative. Hence, we can apply the conditions for the existence of a stable steady state of LCS in Ref. [1] to the biological scenario of cell lineages. In order to use those conditions, let us introduce some mathematical definitions and terminology from Ref. [1].

- If $A$ is an $n \times n$ matrix, $G(A)$ is the *graph* (or *network*) with adjacency matrix $A$, that is, the graph with $n$ nodes and a link from node $i$ to node $j$ weighted by $a_{ij}$, the $(i, j)$-entry of $A$. Such a link only exists if $a_{ij} \neq 0$, and no such link exists if $a_{ij} = 0$.

- We say that nodes $i$ and $j$ are *strongly connected* if there exists a directed path from node $i$ to node $j$ and from node $j$ to node $i$. This is an equivalence relation on the set of nodes. The maximal subsets of strongly connected nodes (the equivalence classes of this equivalence relation) are called the *strongly connected components (SCCs)* of the graph. Moreover, we identify an SCC with the subgraph generated by its vertices (the subgraph containing all the vertices, and all the links between them).

- The network $G(A)$ can be uniquely decomposed into its strongly connected components (subgraphs) $S_1, S_2, ..., S_h$ [3]. Each connected component $S_k$ is a graph with adjacency matrix $A_k$: namely, if $S_k$ has $n_k$ nodes, $A_k$ is the $n_k \times n_k$ submatrix of $A$ consisting of the rows and columns corresponding to the nodes of $S_k$. Each $A_k$ has non-negative off-diagonal elements (because it is a submatrix of $A$), that is, it is a *Metzler matrix*. Moreover, $A_k$ is irreducible (since $S_k$ is strongly connected). Therefore, the Perron-Frobenius theorem ensures that a unique, simple and real maximal eigenvalue $\mu_k$ exists [4]. This is called the *dominant eigenvalue* of $A_k$, and corresponds to the growth parameter defined in the Main Text. We call an SCC $S_k$ *critical* if $\mu_k = 0$, *sub-critical* if $\mu_k < 0$, and *super-critical* if $\mu_k > 0$.

- Given two SCCs, $S_k$ and $S_l$, if there is a (directed) path from a node in $S_k$ to a node in $S_l$, we say that $S_k$ is *upstream* of $S_k$ and that $S_l$ *downstream* of $S_k$. (They are both partial orders in the set of SCCs of $G(A)$.) Note that there cannot be (directed) paths from $S_k$ to $S_l$ *and* from $S_l$ to $S_k$: otherwise, all nodes in both components would be mutually reachable and thus $S_k$ and $S_l$ would merge into a single SCC.

- A *steady state* of the dynamical system $\frac{d}{dt}\boldsymbol{x} = A\boldsymbol{x}$ is a state $\boldsymbol{x}^*$ for which $\frac{d}{dt}\boldsymbol{x}^* = 0$ (i.e. there is no change over time after reaching this state) or, equivalently, $A\boldsymbol{x}^* = 0$. Clearly, the zero vector $\boldsymbol{x}^* = \mathbf{0}$ is a steady state, called the *trivial* steady state. A steady state is called (Lyapunov) *stable* if a small initial deviation from $\boldsymbol{x}^*$ leads to a small deviations $\boldsymbol{x}(t)$ at any time. More accurately: there exists a constant $C > 0$ such that $|\boldsymbol{x}(t) - \boldsymbol{x}^*| < C|\boldsymbol{x}_0 - \boldsymbol{x}^*|$, for all times $t$, where $\boldsymbol{x}_0 = \boldsymbol{x}(t = t_0)$ is the initial condition, sufficiently close to $\boldsymbol{x}^*$.

According to Ref. [1], the following are necessary and sufficient conditions for the existence of a non-trivial (Lyapunov) stable steady state in a LCS.

**Theorem 1** *The LCS $\frac{d}{dt}\boldsymbol{x} = A\boldsymbol{x}$ possesses a non-trivial, Lyapunov stable, steady state if and only if*

1. *G(A) does not contain any super-critical SCC.*
2. *There is at least one critical SCC.*
3. *There are no directed paths from a critical SCC to another. (Equivalently, there are no other critical SCCs downstream, or upstream, of any critical SCC.)*

Furthermore [1], any SCC upstream of a critical SCC (note that this must be a sub-critical SCC) will 'vanish' in the following sense.

**Theorem 2** *Any SCC which is upstream of a critical SCC must be trivial, that is, the stationary state on the nodes of that SCC $S_l$ vanishes, $\boldsymbol{x}^*|_{S_l} = 0$, where $\boldsymbol{x}^*|_{S_l} = (x^*_{i_1}, x^*_{i_2}, ...)|_{i_1,i_2,...\in S_k}$.*

Next, we translate these mathematical statements to the biological context of the Main Text.

- The equation $\frac{d}{dt}\boldsymbol{n}(t) = A\boldsymbol{n}(t)$ in the Main Text (Eq. **[5]** in Box 1) models the dynamics of the mean cell numbers $n_i(t)$ of cell state $i$ at time $t$. This system is a LCS since the transitions rates $\kappa_{i\to j} \geqslant 0$ are non-negative. The graph $G(A)$ is the *cell state network.*
- A directed path in the graph $G(A)$ corresponds to a *cell state trajectory.* The graph $G(A)$ therefore contains all possible cell state trajectories.
- According to the definition in the Main Text, a strongly connected component in $G(A)$, characterised by reversibility of cell state trajectories, corresponds to a *cell type.* Hence, we associate a cell type $T_k$ to each SCC $S_k$ $(k = 1, ...h)$ of $G(A)$.
- *Homeostasis* is characterised by a constant distribution of cell types in a (potentially dynamic) tissue. Mathematically, this corresponds to

a steady state $\boldsymbol{n}^*$ of the LCS above, that is, $\frac{d}{dt}\boldsymbol{n}^* = 0$. In addition, we require that this state is stable and non-trivial ($\boldsymbol{n}^* \neq \boldsymbol{0}$). Since for linear systems the only asymptotically stable state is trivial $\boldsymbol{n}^* = 0$, we use the weaker condition of Lyapunov stability (see above) to define a homeostatic state.

- The dominant eigenvector $\mu_k$ of the adjacency matrix $A_k$ of the SCC $S_k$ is the *growth parameter* of cell type $T_k$. It determines the cell dynamics: if $\mu_k > 0$ then $|\boldsymbol{n}| \to \infty$ (the cell number diverges), if $\mu_k < 0$ then $|\boldsymbol{n}| \to 0$ (the cell number vanishes), and if $\mu_k = 0$ then the mean cell number stays constant over time. We use this behaviour to define cell types: if $\mu_k > 0$, we call the cell type $T_k$ *hyper-proliferative* (corresponding to a super-critical SCC), if $\mu_k < 0$ then $T_k$ is *transient* (corresponding to a sub-critical SCC), and if $\mu_k = 0$ then $T_k$ is a *self-renewing* cell type (corresponding to a critical SCC)[1].

Since Eq. [**5**] in the Main Text (Box 1) is an LCS, Theorems 1 and 2 above apply in this biological context, resulting in the following.

Theorem 1 gives the following necessary and sufficient conditions for homeostasis.

1. There cannot be any hyper-proliferative cell type.
2. There must be at least one self-renewing cell type.
3. Two self-renewing cell types can never be on the same cell trajectory (i.e. connected by a directed path).

Moreover, following from Theorem 2,

4. Any cells of a type upstream of a self-renewing cell type in the lineage hierarchy must vanish in the homeostatic state (they may however be present during development).

We have, in particular, that the following must hold in any homeostatic renewing tissue.

- (From (4)) Self-renewing cell types must always be at the beginning of a homeostatic trajectory, i.e. at the apex of the homeostatic lineage hierarchy. Conversely, each apex of a lineage hierarchy must be a self-renewing cell type. We define those cell types as *adult stem cells.* We note that multiple apices of a lineage hierarchy may exist.

[1] In case of an inert tissue, $\mu_k = 0$ could also denote *inert* cell types, however, throughout this work we only consider renewing tissues, and thus $\mu_k = 0$ must involve proliferative activity. Proliferative activity without net gain or loss of cells is self-renewing.

- When an adult stem cell divides, one or two of its daughter cells may be stem cells of the same type, but not of another stem cell type (otherwise, this would contradict condition 3 above).
- Daughter cells of adult stem cells may not be developing cells, and daughter cells of committed cells may never be adult stem cells nor developing cells (this would contradict condition 3 above).

### 2 Regulation of stemness by crowding feedback

We now consider the situation that cells are able to sense the cell density in their local environment – commonly called the 'niche' – and adjust their dynamic behaviour accordingly (*crowding feedback*). In our model, this means that the kinetic parameters $\lambda_i, \omega_{ij}, d_i$ $(i, j = 1, 2, ..., m)$ depend on the number of cells of a certain type. Let us consider the dynamics of a cell type $T$, which resides at the apex of the lineage hierarchy, and assume that parameters depend only on the number of cells of type $T$, $N = \sum_{i \in T} n_i$ (see also discussion in Main Text). Hence,

$$\frac{d}{dt}\boldsymbol{n}(t) = A(N(t))\,\boldsymbol{n}(t) \qquad (1)$$

where here, $\boldsymbol{n} = (n_1, ..., n_m)$ comprises only the cells of the type $T$ ($m$ is the number of states of type $T$) and $A$ is the sub-matrix for the dynamics of type $T$ according to Eq. [**6**], Main Text[2]. The dependence $A = A(N)$ means that the elements of the matrix $A$ are functions of $N$, and therefore also the dominant eigenvalue of $A$ (the *growth parameter* in the Main Text) is a function of $N$, $\mu = \mu(N(t))$, which thus becomes a dynamic quantity.

In the following we show that under the right conditions, this feedback contributes to the regulation of a cell type's self-renewal capacity: it can render a cell type to become a self-renewing cell type, i.e. $\mu = 0$ according to our definition, even if it is not *a priori* a self-renewing cell type. For a self-renewing state to prevail, there must exist a particular cell configuration $\boldsymbol{n}^*$ with $N^* = \sum_{i \in T} n_i^*$ such that $\mu(N^*) = 0$. Furthermore, the cell configuration in type $T$, $\boldsymbol{n}$, must converge to $\boldsymbol{n}^*$ over time, which is the case if $\boldsymbol{n}^*$ is an asymptotically stable fixed point of the dynamical system, Eq. (1), and if initially $\mu$ is sufficiently close to zero. Hence, the condition that $\boldsymbol{n}^*$ is asymptotically stable state of Eq. (1), implies that cell type $T$ is self-renewing, and if it is the apex of a non-trivial lineage (i.e. with more than one cell type) this means that it will acquire and maintain stemness.

[2]Note that here, for convenience, we changed the notation compared to the Main Text.

To find such conditions, we first transform Eq. (1) into the Jordan basis of $A$, consisting of the linear independent basis vectors $\boldsymbol{e}_1, \boldsymbol{e}_2, ..., \boldsymbol{e}_m$. Of these basis vectors $\boldsymbol{e}_1, \boldsymbol{e}_2, ..., \boldsymbol{e}_q$ ($q \leqslant m$) are normalised eigenvectors of $A$, whereby $\boldsymbol{e}_j$ is eigenvector to the $j$-th eigenvalue $\nu_j$, associated with the $j$-th Jordan block, and $\boldsymbol{e}_{q+1}, .., \boldsymbol{e}_m$ are complementary basis vectors which are not eigenvectors. Note that the latter set of basis vectors is only non-empty if the number of linear independent eigenvectors, $q$, is fewer than the dimension, $m$, of $A$, i.e. when $A$ is not diagonisable; then there are $q < m$ Jordan blocks. When numbering eigenvalues in descending order, $\boldsymbol{e}_1$ is the eigenvector to maximal eigenvalue $\mu = \nu_1 = \max_i\{\nu_i\}$. Since $A$ is a Metzler matrix, the eigenvalue with maximal real part is a dominant eigenvalue, i.e. it is simple, real, and the corresponding Jordan block is scalar [4]. We can therefore write Eq. (1) in the Jordan basis,

$$\frac{d}{dt}\tilde{\boldsymbol{n}}(t) = \tilde{A}(N)\,\tilde{\boldsymbol{n}} \tag{2}$$

where $\tilde{\boldsymbol{n}} = (\tilde{n}_1, \tilde{n}_2, ..., \tilde{n}_m)$ is the representation of vector $\boldsymbol{n} = \sum_j \tilde{n}_j \boldsymbol{e}_j$ in the Jordan basis, and $\tilde{A}(N)$ is the Jordan normal form of the matrix $A(N)$, which can be written as,

$$\tilde{A}(N) = \begin{pmatrix} \mu(N) & 0 \\ 0 & \tilde{A}_{rem}(N) \end{pmatrix}, \tag{3}$$

where the first Jordan block is scalar (a $1 \times 1$ matrix with $\mu(N)$ being its only entry). $\tilde{A}_{rem}$ is a block matrix containing all other Jordan blocks. Crucially, the real values of eigenvalues of $\tilde{A}_{rem}(N)$ are all strictly smaller than $\mu(N)$ for all $N$.

The condition for self-renewal, $\mu(N^*) = 0$ means that $\boldsymbol{e}_1$ is 0-eigenvector, thus $\tilde{A}(N^*)(C\boldsymbol{e}_1) = 0$ for any factor $C \in \mathbb{R}$. Since $N = \sum_{i \in T} n_i = \boldsymbol{1} \cdot \tilde{\boldsymbol{n}}$, where $\boldsymbol{1} = (1, 1, ..., 1)$ in the original basis, the configuration $\tilde{\boldsymbol{n}}^* = (\tilde{n}_1^*, \tilde{n}_2^*, ...) = \frac{N^*}{\boldsymbol{1} \cdot \boldsymbol{e}_1}\boldsymbol{e}_1$ is a steady state of Eq. (2), and satisfies $\mu(N^*) = 0$. This means that $\tilde{n}_1^* = \frac{N^*}{\boldsymbol{1} \cdot \boldsymbol{e}_1}$ while $\tilde{n}_i^* = 0$ for all $i \neq 1$.

For assessing the stability of $\boldsymbol{n}^*$, we consider the Jacobian matrix,

$$\tilde{J} = [\tilde{J}_{ij}] \text{ with } \tilde{J}_{ij} = \frac{\partial(\frac{d}{dt}\tilde{n}_i)}{\partial \tilde{n}_j} = \frac{\partial(\tilde{A}\boldsymbol{n})_i}{\partial \tilde{n}_j} = \tilde{a}_{ij} + \sum_k \frac{\partial \tilde{a}_{ik}}{\partial \tilde{n}_j}\tilde{n}_k \tag{4}$$

where $a_{ij}$ is the $i, j$-element of $A$. Thus the elements Jacobian matrix at the steady state $\boldsymbol{n}^* = (\frac{N^*}{\boldsymbol{1} \cdot \boldsymbol{e}_1}, 0, ..., 0)$ are

$$\tilde{J}_{ij} = \tilde{a}_{ij} + \frac{\partial \tilde{a}_{i1}(N^*)}{\partial N}N^* \tag{5}$$

where we used that $\frac{\partial \tilde{a}_{ij}}{\partial \tilde{n}_1} = \frac{\partial \tilde{a}_{ij}}{\partial N}\frac{\partial N}{\partial \tilde{n}_1} = \frac{\partial \tilde{a}_{ij}}{\partial N}(\mathbf{1}\cdot \boldsymbol{e}_1)$. This can be written as

$$\tilde{J} = \begin{pmatrix} \overbrace{\mu(N^*)}^{=0} + \frac{\partial \mu(N^*)}{\partial N}N^* & \frac{\partial \mu(N^*)}{\partial N}N^* \\ 0 & A_{\text{rem}} \end{pmatrix} \tag{6}$$

since all $a_{i1} = 0$ for $i > 1$ and $a_{1j} = 0$ for $j > 1$. Now, the condition that $\boldsymbol{n}^*$ is asymptotically stable is that all eigenvalues of $J$ (and thus of $\tilde{J}$) are negative [5]. Since $\tilde{J}$ is in an upper triangular block form, the eigenvalues are simply the union of the eigenvalues of $\tilde{A}_{rem}$ and $\mu(N^*) + \frac{\partial \mu(N^*)}{\partial N}\tilde{N}^* = \frac{\partial \mu(N^*)}{\partial N}N^*$. And since the eigenvalues of $\tilde{A}_{rem}$ are all smaller than $\mu(N^*) = 0$, they are all negative and thus the condition for an asymptotically stable steady state is that the remaining eigenvalue, $\frac{\partial \mu(N^*)}{\partial N}N^*$ is negative. Since $N^*$ is positive, this is fulfilled (sufficient condition[3]) if

$$\frac{\partial \mu}{\partial N} = \sum_{ij} \frac{\partial \mu}{\partial a_{ij}}\frac{\partial a_{ij}}{\partial N} < 0 \tag{7}$$

By representing the matrix entries $a_{ij}$ in terms of the system parameters $\alpha_l$, where $\alpha_l$ ($l = 1, ..., m^2 + m$) can be any of the parameters $\lambda_i, \omega_{ij}, d_i$, $i, j = 1, 2, ..., m$, we get the sufficient condition for (asymptotically stable) homeostasis,

$$0 > \sum_{ijl} \frac{\partial \mu}{\partial a_{ij}}\frac{\partial a_{ij}}{\partial \alpha_l}\frac{\partial \alpha_l}{\partial N} = \sum_{l} \frac{\partial \mu}{\partial \alpha_l}\frac{\partial \alpha_l}{\partial N} \tag{8}$$

which is the condition, Eq. [**10**], given in the Main Text.

Condition (8) assures that $\boldsymbol{n}$ converges to $\boldsymbol{n}^*$, and thus $\mu(N)$ converges to $\mu(N^*) = 0$ if the initial conditions are sufficiently close to $\mu = 0$. However, the condition that $\frac{d\mu}{dN} < N$ for all $N$ assures at the same time that only one steady state, namely $N^*$, can exist. Since this is an asymptotically stable fixed point, all initial condition must converge to it.

Hence, the condition (8) represents a sufficient condition that $\mu$ converges to $\mu = 0$, i.e. cell type $T$ attains the self-renewal property. If that cell type $T$ is at the apex of a lineage hierarchy this means that $T$ is a stem cell type. From condition (8) also follows a sufficient sign condition for homeostasis: if all terms $\frac{\partial \mu}{\partial \alpha_l}\frac{\partial \alpha_l}{\partial N}$ are negative, which is fulfilled if the sign of $\frac{\partial \alpha_l}{\partial N}$ is opposite the sign of $\frac{\partial \mu}{\partial \alpha_l}$ for all *relevant* parameters $\alpha_l$ (those parameters with $\frac{\partial \mu}{\partial \alpha_l} \neq 0$),

[3]Note that here we used a stricter sufficient but not essential condition, namely that $\frac{\partial \mu}{\partial N} < 0$ for all $N$. This is required further below.

then condition (8) is fulfilled. Hence, a self-renewing state is assured if all relevant parameters have feedback with the sign of $\frac{\partial \alpha_l}{\partial N}$ opposite to the sign of $\frac{\partial \mu}{\partial \alpha_l}$. This is the sign condition of Box 3 in the Main Text.

To demonstrate how a cell type becomes self-renewing by crowding feedback, we study an example scenario numerically in Figure 1. The figure shows how a particular cell type $T_a$, implemented as a cell state network forming a single SCC (depicted in the top panel), attains a self-renewing state if the sign condition for crowding feedback (Eq. (8) and Box 3, Main Text) is fulfilled. It shows that for various initial conditions which are initially not self-renewing, the growth parameter approaches the value $\mu_a = 0$ over time (right panel), and the cell number $N_a$ becomes stationary (left panel), corresponding to a homeostatic state.

In order to fulfill condition (8) for crowding feedback in that example, we chose each kinetic parameter $\alpha_i \in \{\lambda_j, d_j, \omega_{jk}\}_{j,k=1,\dots,m}$ being a function of $N_a$. To find suitable functions, we first implemented a corresponding model with constant kinetic parameters $\alpha_i$ (no crowding feedback) and employed a genetic search algorithm [6] to find a set of parameters $\alpha_i^*$ for which the dominant eigenvalue of $A$, $\mu_a$, becomes zero. Then we tested, by slightly varying each parameter $\alpha$ around $\alpha_i^*$, whether $\mu_a$ decreases or increases with $\alpha_i$, to determine the sign of $\frac{d\mu_a}{d\alpha_i}$. If $\frac{d\mu_a}{d\alpha_i} > 0$, we choose $\alpha_i(N_a) = K_i/N_a$ so that $\frac{\partial \alpha_i}{\partial N_a} < 0$, and if $\frac{d\mu_a}{d\alpha} < 0$ we choose $\alpha = K_i N_a$ so that $\frac{\partial \alpha_i}{\partial N_a} > 0$, to fulfill the the sign condition (and thus condition (8)). The dependence parameter $K_i$ is chosen for each kinetic parameter $\alpha_i$ as $K_i = 1000\alpha_i^*$, if $\frac{d\mu_a}{d\alpha_i} > 0$, and $K_i = \alpha_i^*/1000$, if $\frac{d\mu_a}{d\alpha} < 0$, respectively, so that the target stationary cell number is $N_a^* = 1000$. We then solved the differential equations describing this system, Eq. (1), numerically for three sets of initial conditions, '$H'$ : $N(t=0) = N_a^*$, '$P_1'$ : $N(t=0) = 1.2N_a^*$ and '$P_2'$ : $N(t=0) = 0.8N_a^*$, and ploted $N_a(t)$ (left panel) and $\mu_a(t)$ (right panel). The conditions $P_1$ and $P_2$ correspond to situations when cell type $T_a$ is initially not self-renewing, visible in the right panel as $\mu_a(t=0) \neq 0$. We observe that for either initial condition the system eventually attains a steady state $N_a \to N_a^*$ (left panel) and self-renewal property $\mu_a \to 0$ (right panel) over time $t$.

### References

[1] P. Greulich, B. D. MacArthur, C. Parigini, and R. J. Sánchez García. Stability and steady state of complex cooperative systems: a diakoptic approach. *Royal Society Open Science*, 6:191090, 2019.

[2] M. W. Hirsch and H. Smith. Monotone dynamical systems. *Handbook of differential equations: Ordinary Differential Equations*, 2:57, 2006.

[3] B. Bollobás. *Modern graph theory.* Springer Science & Business Media, 2013.

[4] K. J. Arrow. A "dynamic" proof of the Frobenuis-Perron theorem for Metzler matrices. In *Probability, Statistics, and Mathematics*, pp. 17, 1989.

[5] S. H. Strogatz. *Nonlinear Dynamics and Chaos: With Applications to Physics, Biology, Chemistry, and Engineering.* CRC Press, 1994.

[6] D. E. Goldberg. *Genetic Algorithms in Search, Optimization and Machine Learning.* Addison-Wesley Longman Publishing Co., 1st edition, 1989.

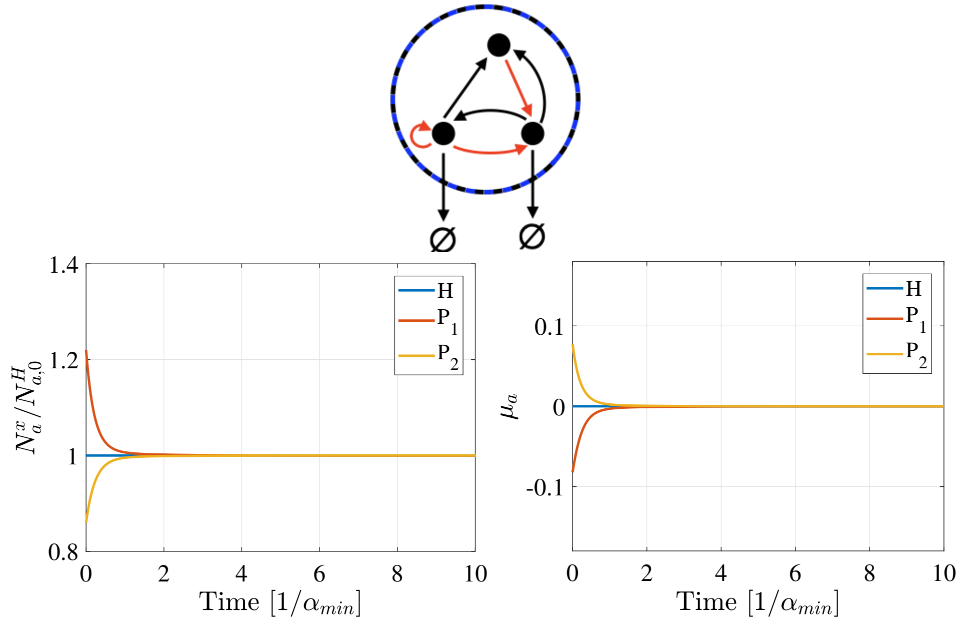


Figure 1: Example of crowding-dependent regulation of stemness for a cell state network with 3 states (shown on the top, with symbols and colours as in Fig. 3 of the Main Text). For each kinetic parameter $\alpha$ we chose the crowding dependence such that the criterion, Eq. (8), is fulfilled, namely $\alpha(N_a) = K/N_a$ if $d\mu_a/d\alpha > 0$, and $\alpha(N_a) = KN_a$; if $d\mu_a/d\alpha < 0$ (see text for how the sign of $d\mu_a/d\alpha$ is found). The left panel shows the time evolution of mean cell numbers $N_a$ (obtained as numerical solutions of the dynamic equation Eq. (1) for type $T_a$), for different initial sets of kinetic parameters, written as $H$, $P_1$ and $P_2$. The right panel similarly shows the time evolution of the growth parameter $\mu_a$. Here $H$ denotes a choice of parameters such that type $T_a$ is initially a self-renewing type with $N_a(t=0) = N_a^*$ (the tissue is in "homeostasis"), while for $P_1$ $N_a(t=0) = 1.2N_a^*$ and for $P_2$, $N_a(t = 0) = 0.8N_a^*$, resulting in initial parameters that render $T_a$ not self-renewing. Time is scaled by the inverse of the smallest rate $\alpha_{min} = \min_{i,j}\{\lambda_i, \omega_{i\to j}, d_i\}$ at $N_a = N_{a,0}^H = N_a^*$. Crucially, we observe that even if $T_a$ is not initially self-renewing ($\mu_a(t=0) \neq 0$), i.e. for parameter choices $P_{1,2}$, the parameters self-adjust over time such that the growth parameter attains $\mu_a = 0$ (right panel). This is also reflected in the the time evolution of cell numbers, which become stationary, $N_a \to N_{a,0}^H = N_a^*$ (left panel).
